# Demonstration of re-epithelialization in a bioprinted human skin equivalent wound model

**DOI:** 10.1101/2020.06.15.152140

**Authors:** Carlos Poblete Jara, Carolina Motter Catarino, Yuguo Lei, Lício Augusto Velloso, Pankaj Karande, William H. Velander, Eliana Pereira de Araujo

## Abstract

**Objective:** The development of an in vitro platform for modeling human skin injury and the re-epithelization process.

**Approach:** A fibrin provisional matrix (FPM) was installed into a wound facsimile of a bioprinted human skin equivalent (HSE). A mixture of plasma-derived fibrinogen-containing factor XIII, fibronectin, thrombin, and macrophages (an FPM “bioink”) was extruded into the wound site. The surrounding *in vitro* tissue culture became a source of keratinocytes to achieve wound closure by a re-epithelialization process signaled by the FPM.

**Results:** An *in vitro* analog of wound closure and re-epithelialization by keratinocytes occurred over the FPM after a normal migration initiation at 3 days.

**Innovation:** A physiologic mixture of macrophage/fibrinogen/fibronectin that supports macrophage differentiation was applied to a mechanically wounded, bioprinted dermal tissue. We developed a transitional culture medium to mimic the changing microenvironment during the initial phases of wound healing. As a reference, we temporally compared our *in vitro* model with a murine skin wound healing.

**Conclusion:** This co-culture model was shown to temporally synchronize a re-epithelization process for initiation of keratinocyte migration from a surrounding tissue and the migration process over the top of an FPM. A future study of the analogous subepithelial healing pathway is envisioned using the same *in vitro* bioprinted tissue study platform for co-culture of keratinocytes, melanocytes, fibroblasts, endothelial cells, and macrophages using more specialized FPMs.

## Introduction

Cutaneous healing begins with a hemostatic response that integrates several physiologic processes: the cessation of blood loss, immediate wound stabilization by sealing tissue from the environment, the initiation of the inflammatory cell response (1), and the simultaneous installation of the fibrin provisional matrix (FPM) to initiate the recruitment and proliferation of healing cells. Thus, the FPM mediates the initiation of re-epithelialization and the construction of a well vascularized, provisional subepithelial tissue. The FPM is penultimately generated by the activation of thrombin from the coagulation cascade and by the thrombin-activation of coagulation factor XIII (XIIIa).

The re-establishment of the epidermis is a fibrin-receptor-mediated process that guides keratinocyte migration from the interfollicular epidermis that surrounds the FPM sealed wound surface. While keratinocytes migrate along the FPM to the wound center, they are continually re-supplied by proliferating keratinocytes in order to complete the wound closure process (1, 2). Provisional subdermal tissue repair is simultaneously initiated to re-epithelialization by the infiltration of the FPM by fibroblasts, endothelial cells and macrophages (3, 4).

*In vitro* wound healing models are advancing to 3D culture formats: these models strive to emulate cell interactions with clustered integrins positioned all around the cell membrane surface. However, engineering of the FPM-cell interface with a complement of different cell types does not re-capitulate the natural tissue healing process. This study expands the *in vitro* 3D skin equivalent co-culture platform to include the installation of a physiologically relevant FPM. This includes using the physiological ratio of macrophages to fibrin surface area (5), the physiological concentration of fibrinogen used to form fibrin (6), and the spatial positioning of the re-epithelialization behavior of keratinocytes over FPM at the skin insult (1). We include a physiological concentration of human fibrinogen and plasma Fibronectin (pFN) (6) and macrophages during the formation of fibrin. The resultant FPM simulates what typically exists during the normal initiation of granulation tissue construction beginning at about 24 hours after injury (5). This FPM fills the subepithelial wound space and within a surrounding store of keratinocytes derived from dermal tissue after a wound insult (1). Our study presents the re-epithelialization and corresponding wound closure phenomena stimulated within a bio-printed tissue co-culture platform. A temporal comparison is made between dermal wound healing in the 3D bio-printed platform and a murine dermal wound model. This platform also serves as a basis for future granulation tissue formation studies.

### The Pre-Clinical Utility of the 3D Bioprinting Wound Healing Platform

Annually, millions of animals are used for experimental purposes (7) looking for new and better treatments for wound management. Small animals and pigs models have been used to study healing of cutaneous wounds (8, 9). However, the outcomes from these experiments cannot be applied directly to humans due to species-specific differences (10). In addition to the difficulty of transferring the findings, the model of full thickness wounds in experimental animals (11) is limited by ethical constraints. In 1959, William Russell and Rex Burch proposed in the “The Principles of Human Experimental Technique” where possible animals should be replaced with alternatives that minimize animal numbers while preventing pain and distress (12, 13). The 3D human skin equivalent (HSE) studied has utility to reduce the role of animal testing in preclinical studies.

## Materials and Methods

### Experimental animals

Eight-week-old male C57BL/6J isogenic mice (n=5) were obtained from the Breeding Animal Center of University of Campinas. Animals were maintained under pathogen-free conditions in individual cages on a 12:12 hour dark:light cycle at 21–23°C. Mice received chow and water ad libitum. Mice were anesthetized with intraperitoneal injections (according to body weight) using ketamine hydrochloride 80 mg/kg and xylazine chlorhydrate 8 mg/kg. Under general anesthesia, we created two full-thickness excisional wounds (6.0 mm diameter each) on the back of each animal, according to the mouse excisional wound splinted model protocol (11). Animal experiments were approved by The Animal Ethical Committee at the University of Campinas, Brazil (certificate of approval no. 4467-1).

### Animal photo documentation

Photo documentation of the wound healing process was obtained using a D610 Nikon digital camera (Nikon Systems, Inc., Tokyo, Japan). To ensure a similar distance from the camera to wound site, we used a stand and the same person took the photos.

### Fibrin Provisional Matrix (FPM)–macrophage and FPM– Human Endothelial Cells (HUVEC) bioink mixture formulations

We created two master solutions to produce a fibrin sealant delivering pFN and macrophages to the mechanical insult of the tissue culture. Mixture 1 contained whole population F1 (2.5 mg/ml) from normal human plasma, pFN (0.25 mg/ml) and 200 macrophages/μL or 500 HUVEC/μL in phosphate buffered saline (PBS) 1× suspension. Polymerization and protein crosslinking were catalyzed by Mixture 2, which contained thrombin (5 U/ml), calcium (5 mM), and FXIII (50 μg/ml). The same amount of PBS 1× without cells was used as the control. To convert fibrinogen to fibrin, we used F2 (activated thrombin). All values are the final concentration in the fibrin polymer.

### FPM droplet studies for bio-ink formulation

FPM–macrophage mixture droplets (30 μL and 50 μL) were applied to autoclaved glass slides with 8-well silicone cell culture chamber wells (Ibidi) by first combining Mixture 1 and Mixture 2, with 200 macrophages/μL or 500 HUVEC/μL (**Figure 1a**). After 15 minutes at 37°C in a 7% CO_2_ incubator, the polymerized/cross-linked FPM, 400 μL Iscove's Modified Dulbecco's Medium (IMDM) 20% Fetal Bovine Serum (FBS) medium was added to each well and cultured at 37°C in a 7% CO_2_ incubator for 7 days. The m acroscopic F PM stability was e valuated u sing photographs t aken 0 a nd 7 days a fter confection using a 1× Olympus SZX16 microscope and cellSens Standars Software 1.11 (Olympus Corporation) with KL1500 LCD (Schott) 3200K top illumination.

**Figure 1.**
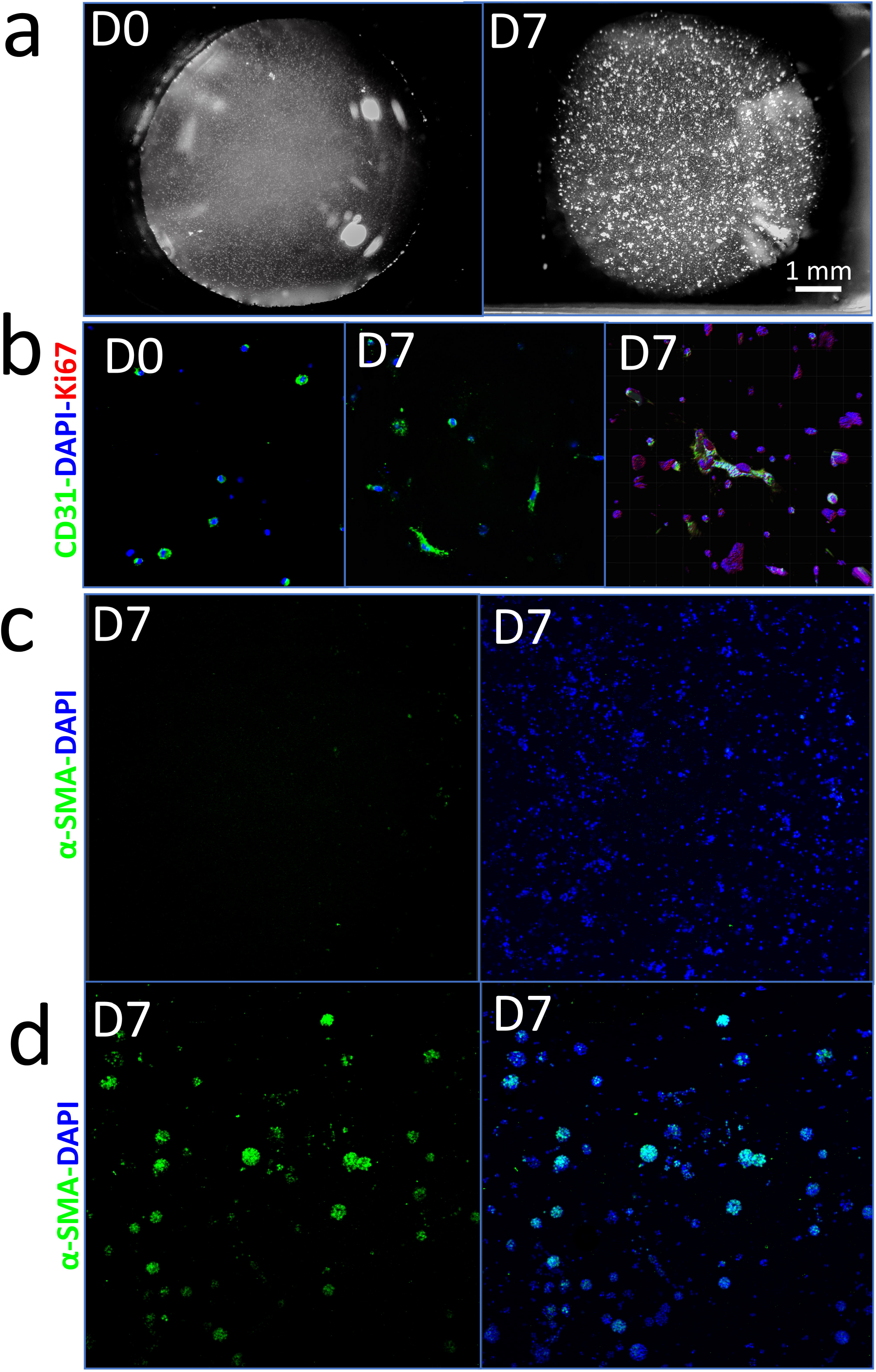
Droplet experiment using the fibrin clot bioink. Day 0 and day 7 after the droplet experiment (a). CD31, DAPI, and Ki67 labeling using HUVECs in the fibrin clot on days 0 and 7 (b). a-SMA and DAPI labeling using human macrophages in the fibrin clot structure (c–d), using inflammatory medium and proliferative medium (c), and using creation medium (d).

### Primary cell culture preparations

Primary keratinocytes and fibroblasts were harvested and isolated from human skin, as previously described (14), and under institutional review board approved #AAAB2666 protocol. About 2–4 passages of primary keratinocytes were made in culture in 37°C in a 7 % CO_2_ incubator with KBM-Gold supplemented with KGM-Gold Single Quots (Lonza) medium to 80% confluence in 100 mm Petri dishes. Primary fibroblast passages 2–6 were achieved in culture in 37°C in a 5% CO_2_ incubator with DMEM High Glucose (Gibco) with 10% FBS to 80% confluence in 100 mm Petri dishes. HUVEC cells were cultured to <8 passages in EBM-2 (Lonza) supplemented with EGM-2 Single Quots (Lonza). Bone marrow human macrophage (KG-1 ATCC^®^ CCL246™) passages 2–5 were cultured in cell suspension flasks using IMDM medium (Gibco™ GlutaMAX™, HEPES) with 20% FBS. Macrophage suspensions were cultured at 37°C in a 5% CO_2_ incubator. We maintained macrophages at a concentration of less than 1 × 10^6^ viable cells/mL, replacing the culture m edium twice a week by centrifugation at 130 × g for 7 minutes.

### Preparation of printed human skin equivalent (HSE)

Human skin equivalent (HSE) was prepared according to previous protocols (14). In brief Collagen I (2.5 mg/ml final solution), reconstitution buffer 10× (**Table 1**), 1:1 HAM:F12 10X, and primary fibroblast (1.5 × 10^5^ final solution) were mixed, gently homogenized, and then manipulated over a cooled metal rack at 4°C. The solution was manipulated over a cooler cartridge head to 4–8°C, and then applied to a printer bed at 4°C with 2200 μL of the mixture into a 3 mL-printer cartridge. This collagen–fibroblast solution was extruded (**Table 2**) into each of the 6 wells of a 24 mm transwell (200 s of extrusion; pressure, 50 kpa; Nozzle type, 30G; no top piston). Next, more 1100 μL (130 s of extrusion; pressure, 50 kpa; Nozzle type, 30G; no top piston) were mixtured in the previous transwell totaling 3000 μL per transwell (**Figure 2a**). Then, 6-well plates were incubated in a 5% CO_2_ incubator for 30 minutes at 37°C to assure gel formation. Next, 0.5 ml of DMEM was applied to the top of each individual culturing skin equivalent, with 2 ml between the transwell and plate, while incubating for 1 hour at 37°C in the 5% CO_2_.

**Table 1.**
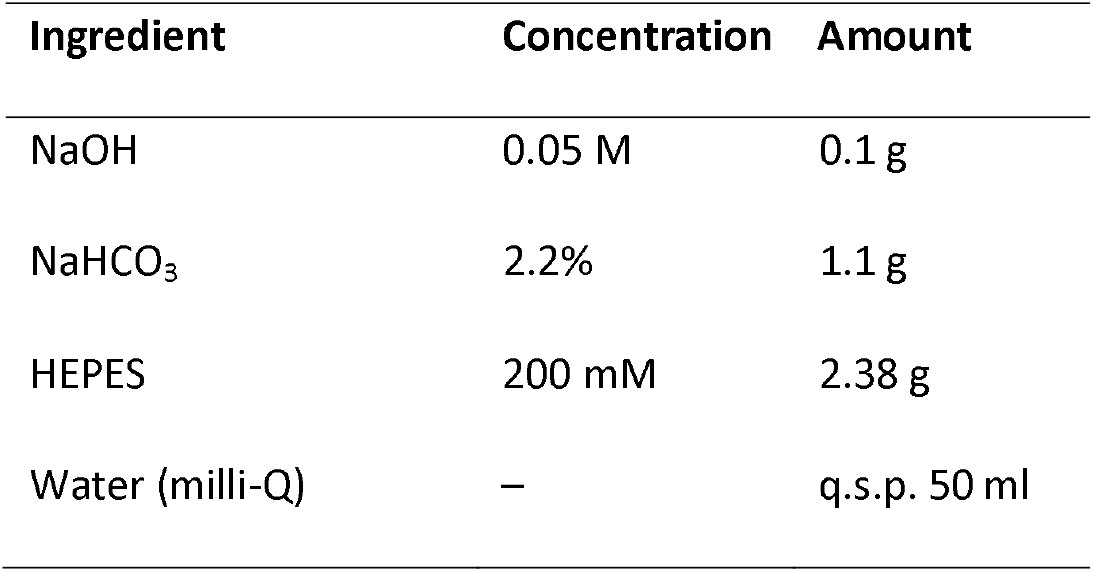
Reconstitution buffer.

| Ingredient | Concentration | Amount |
| --- | --- | --- |
| NaOH | 0.05 M | 0.1 g |
| NaHCO<sub>3</sub> | 2.2% | 1.1 g |
| HEPES | 200 mM | 2.38 g |
| Water (milli-Q) | – | q.s.p. 50 ml |

**Table 2.** GCodes for bioprint experiments.

| Dermis | Dermis | Epidermis | Fibrin-clot |
| --- | --- | --- | --- |
| G0 X0 Y0 | G0 X0 Y0 | G0 X0 Y0 | G0 X0 Y0 |
| G0 Z15 | G0 Z15 | G0 Z15 | G0 Z15 |
| T1 | T1 | T1 | T1 |
| G1 Z0 F600 | G1 Z0 F600 | G1 Z1 F600 | G1 Z1 F600 |
| M400 | M400 | M400 | M400 |
| M750 T1 | M750 T1 | M750 T1 | M750 T1 |
| G4 S200.000 | G4 S130.000 | G4 S55.000 | G4 S6.000 |
| G1 Z10 F90 | G1 Z10 F90 | G1 Z10 F90 | G1 Z10 F90 |
| M400 | M400 | M400 | M400 |
| M751 T1 | M751 T1 | M751 T1 | M751 T1 |
| G0 Z15 | G0 Z15 | G0 Z15 | G0 Z15 |
| G1 X0 Y0 | G1 X0 Y0 | G1 X0 Y0 | G1 X0 Y0 |

**Figure 2.**
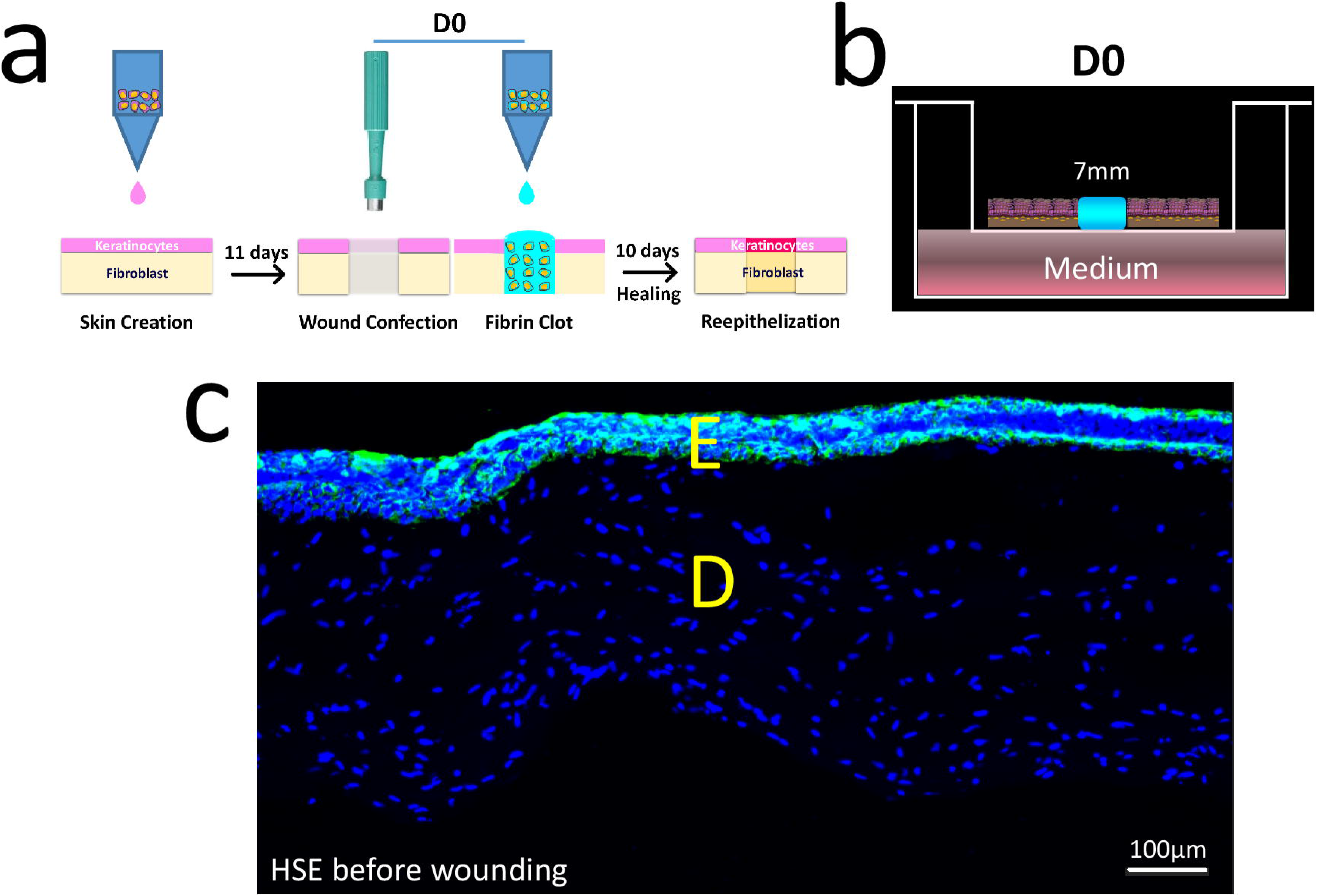
Bioprinted human skin equivalent. Schematic representation of wound creation, wound confection, fibrin clot extrusion, and re-epithelization (a). Human skin equivalent in the air–liquid interphase (b). Cytokeratin-DAPI labeling in a skin equivalent after 10 days of air–liquid interphase (c).

After all DMEM medium was removed, 450 μL of 1:1 KGM gold/Creation medium containing 1 × 10^6^ primary keratinocyte was extruded into the center of each transwell (55 s of extrusion time at 10 kPa pressure using Nozzle type 30G with no top piston). An additional 2 ml of 1:1 KGM gold/Creation medium was added underneath the transwell. Keratinocyte culturing on the top of each skin equivalent layer was maintained for 24 hours to achieve cell adhesion while at 37°C in 7.5% CO_2_. After 24 hours, the keratinocyte media was discarded and each skin equivalent was detached from the transwell walls using a sterile surgical spatula and transferred to the air–liquid interphase, and the apparatus was placed in a deep 6-well plate using sterile forceps. Twelve milliliters of Creation Medium (**Table 3**) was added below each transwell, while avoiding bubble formation where these air–liquid interphase incubation samples were maintained at 37°C in 5% CO_2_. The Culture medium schedule (**Table 3**) was maintained for 10 days.

**Table 3.**
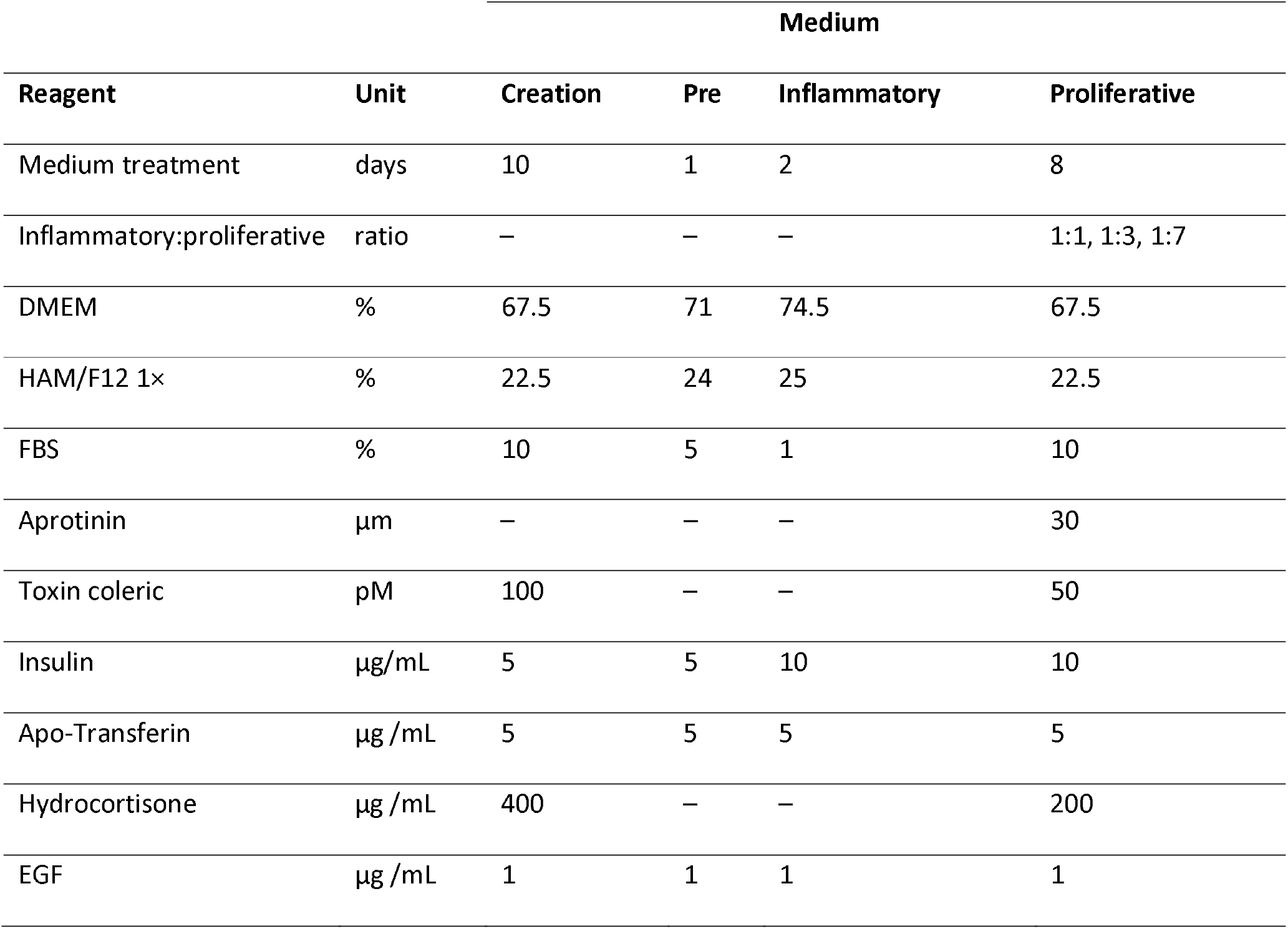
Culture medium schedule.

A surplus stock of 10% solutions for treating all cell layers is used for bubble formation management. The sterility of the print chamber was maintained by UV light within a HEPA-filtered chamber. The initial extrusion of collagen–fibroblast solutions used a Z0 position 1 mm above the transwell membrane, with secondary extrusions (set Z0 position=1mm application above the previous dermal solution layer). Keratinocyte solution extrusion used a Z0 position of 1 mm, applied above the polymerized collagen–fibroblast gel layer.

### Wound creation of the HSE

Full-thickness injuries were made at the middle of each HSE sample on day 11 using a sterile 4 mm or 7 mm dermal biopsy punch (15). Each injured HSE was placed in a dry Petri dish after treatment with 10 ml of 1:1 Trypsin inhibitor and HEPES for 1 minute to remove traces of culture medium. The injury motion consisted of a single, quick perpendicular stroke accompanied by one rotatory movement to free the punch from the HSE (**Figure 2a, 3b**). The cleared wound HSE cavity was immediately filled with FPM–macrophage “bioink.”

### HSE wound healing photo documentation

The healing process was evaluated through photographs taken 0, 2, 6, 8, and 10 days after wounding using a 1× Olympus SZX16 microscope and cellSens Standard Software 1.11 (Olympus Corporation) with KL1500 LCD (Schott) 3200K top illumination. The wounded areas and re-epithelialized areas were analyzed with ImageJ Software (1.49v).

### Printing fibrin clots in a wounded HSE

Macrophages were washed by centrifugation and suspended in 1× PBS without calcium to eliminate medium traces. The fibrin clot–macrophage bioink containing Mix 1 (2.5 mg/ml F1, 0.25 mg/ml FN, 200 macrophages/μL, and 25 μM aprotinin) was pipetted into a bioprinter cartridge. Forty microliters of fibrin clot bioink was extruded (6 s of extrusion; 10 kpa; 30G needle; 20 mm/s; no top piston; 400 μL of Mixture 1 inside the cartridge) into the center of each wound. Ten microliters of Mixture 2 was manually applied on the top of each fibrin clot bioink (**Figure 2a, b**). The cartridge head and printer bed were maintained at room temperature. We added aprotinin because, in a pilot experiment, the fibrin clot bioink was enzymatically digested during the first 24 hours after wounding the skin (data no shown). All values are final concentration in the fibrin clot bioink. To extrude the fibrin clot bioink, we set the Z0 position to 1 mm above the transwell membrane, in the center of the injured HSE. This extrusion system can be easily replaced by a less restrictive method, such as the use of pipettes.

### HSE culture medium refreshment for managing the transition between the inflammatory phase and proliferative phase cell culture environment

Newly prepared HSEs were maintained for 11 days before injury using “Creation medium.” On day 10 of the pre-wounding phase, the HSE media was refreshed with “Pre-Wound Media” to clear traces of hydrocortisone and toxic choleric. The newly wounded HSE was then maintained for an additional 10 days using a sequence of media designed to mimic the natural transition from the inflammatory phase to the proliferative Phase. The composition of the media regimen is detailed in **Table 3**. After injury, the HSE were refreshed with inflammatory phase media. On day 2 after wounding, 20 μg/ml (final medium concentration) of aprotinin (16) was added to manage FPM degradation. On day 3 post-injury, 50% of the inflammatory phase media (6 ml) was removed and replaced with an equal volume (6 ml) of proliferative phase medium to begin the transition to the proliferative phase (4, 5, 17). This gradual dilution by proliferative phase medium occurred on day 6 and 9, with harvest occurring on day 10.

### Immunofluorescence viability analysis of FPM–macrophage droplet bioink

To assess viability after 7 days in culture, IMDM medium was discarded from droplets of FPM–macrophage. These droplets were then fixed using 4% PFA for 15 minutes (7.5 pH; corrected with NaOH). PFA was discarded and fibrin clots were washed 3 × 15 minutes with PBS 1×. Fibrin clots were blocked for 1 h with PBS 1×, 0.05% Tween 20, and 5% BSA. Antibodies were prepared with PBS 1×, 0.05% Tween 20, and 1% BSA solution. Anti-CD31 (1:50, ab28364) and Ki-67 (1:100, BD 610968) were maintained at 4°C overnight. Secondary antibodies, Alexa 488 (1:200, ab150077) and Alexa 647 (1:200, ab150115), were maintained at 4C° overnight. Vectashield with DAPI was used for mounting and counter nuclear staining.

### Immunofluorescence analysis of HSE cell viability

After 10 days post wounding, the proliferative medium was discarded from the HSE, and these samples were fixed by 4% PFA (7.5 pH; corrected with NaOH), with incubation at 4°C overnight. The PFA was discarded from the samples, followed by three 15-minute washes with PBS 1×. Samples were then embedded in Optimal Cutting Temperature (OCT) inside of a quadrangular prism foil form and quickly frozen using 100% ethanol in dry ice. Tissue sections (50 μM) were obtained using Microm HM505E Cryostat and placed in a glass slide. To remove OCT, tissue sections were washed for 30 minutes in distillated water, and peripheral marking was made using a hydrophobic marker. These tissue sections were blocked for 3 hours with PBS 1×, 0.05% Tween 20, and 5% BSA. Anti-K14 (1:200, ab7800) antibodies in PBS 1×, 0.05% Tween 20, and 1% BSA solution were incubated with samples at 4°C overnight and then warmed for 30 minutes at room temperature. Secondary antibody Anti-Mouse Alexa Fluor 488 (1:1000 Invitrogen A1101) was incubated with samples at room temperature for 4 hours. Vectashield with DAPI was then applied for counter nuclear staining, with images obtained by confocal microscopy (Zeiss LSM 510 Meta Spectral Confocal) at a penetration thickness of 20 μM.

### Hematoxilin/Eosin (HE) staining for histological imaging

After 10 days post wounding, HSE samples were fixed as above. Tissue sections were stained as follows using HE: 30-minute soak in distillated water followed by a 2-minute soak in hematoxylin (Sigma-Aldrich) and rinsing with tap water for 5 minutes. Tissue sections were stained with Eosin (Sigma-Aldrich) soak for 10 minutes, then the following ethanol drying sequence of 5 minute soaks: 50% ethanol soak, 70% ethanol soak, 90% ethanol soak, 95% ethanol soak, 100% ethanol soak, fresh 100% ethanol, fresh 100% ethanol, Xylol I, and fresh Xylol II for the last 5 minutes. Slide assemblies were made with DPX mounting solution and a coverslip.

### Full view tissue image reconstruction

To improve the accuracy of the analysis, we reconstructed the whole skin section (unwounded and injured sample). We used a Nikon digital camera (Nikon Systems, Inc., Tokyo, Japan) to take 10–20 adjacent microphotographs (with 10–20% of common areas; 10× objective) used to reconstruct every skin section. We used the Photomerge tool with the perspective layout in Adobe Photoshop (19.1v).

## Results

### Fibrin Stability and Cell viability of the FPM–cell mixtures

Bioinks made containing Macrophages or HUVECs were observed for 7 days. The macroscopic matrix stability of the FPM–macrophage and FPM–HUVEC bioinks were stable for > 7 days as judged by the lack of macroscopic matrix degradation (**Figure 1a**). The Macrophages or HUVECs cultured withing this FPM droplets were observed to proliferate for the entire 7 day culture period (**Figure 1b**).

### Re-epithelization over an injured HSE sealed with the FPM–macrophage bioink tissue sealant

The entire 21-day HSE and HSE wound healing sequence is schematically shown in **Figure 2a**. HSEs maintained for 1 day while submerged and for 10 days in the air–liquid interphase were harvested and histologically visualized. The keratinocyte layer was observed on top of the fibroblast–collagen layer (**Figure 2c**). A typical unwounded HSE is shown in **Figure 3b** (the left panel shows two different wound sizes that were made using a 4 mm [Figure 3b, center] and a 7 mm [Figure 3b, right] biopsy punch). The wounds were sealed with the FPM–macrophage bioink after injury. After 8 days post wounding, the 4 mm injured HSE showed complete reepithelization (**Figure 3a, d**). After 10 days post wounding, the 7 mm injured area showed complete reepithelization (**Figure 3a, c**). This contrasted the wounded HSE without the FPM–macrophage bioink treatment, which showed an inhibited wound closure (**Figure 3a**). Both 4 mm and 7 mm (**Figure 3c, d**) wounds were observed to mimic the macroscopic animal wound closure (**Figure 3a, e**).

**Figure 3.**
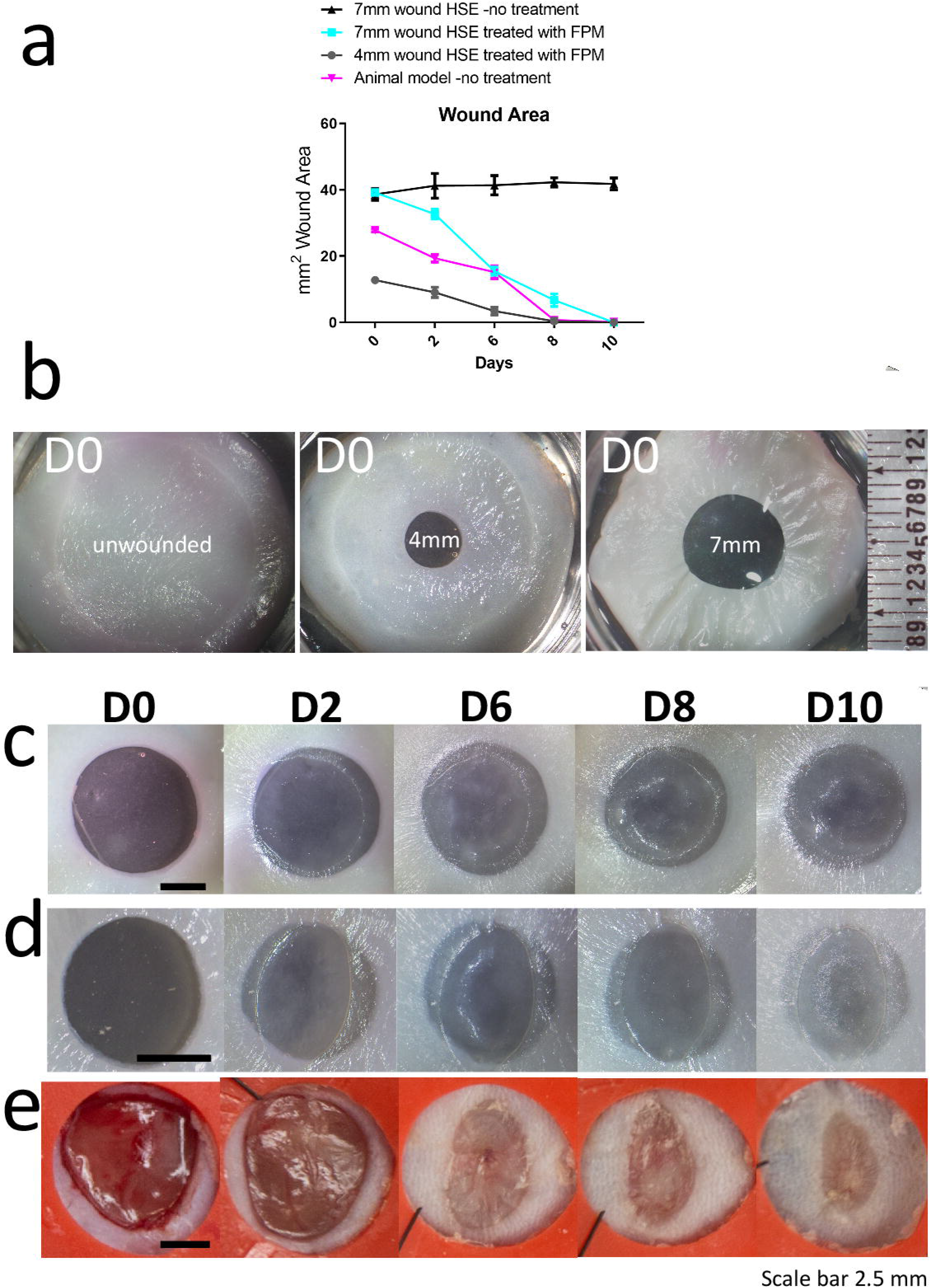
Comparison of the wound healing process with human bioprinted skin and skin from an animal model. Human skin equivalents (n:6) versus animal healing (n:6) rate comparison graph (a). Human skin equivalent 11 days post-creation (D0), unwounded, using a 4 mm or 7 mm biopsy punch (b). *in vitro* wound healing process (c–d) compared to the animal healing process (e) using 7 mm (c), 4 mm (d), and 6 mm (e) biopsy punches. The images are representative of three distinct experiments.

During mammalian skin wound healing, keratinocytes proliferate and migrate from the interfollicular epidermis to the wound center (1, 2). In our model, keratinocytes migrated from the wound edge to the wound center using the fibrin clot–macrophage bioink as a platform (**Figure 4c, d**). In the same way as the animal model (**Figure 4a**), wounded HSE model (**Figure 4b**) regenerated a new layer in the wound center. The re-epithelization in the wounded HSE model is dependent on the initial wound size and the presence of hemostatic fibrin clot bioink (**Figure 3a**).

**Figure 4.**
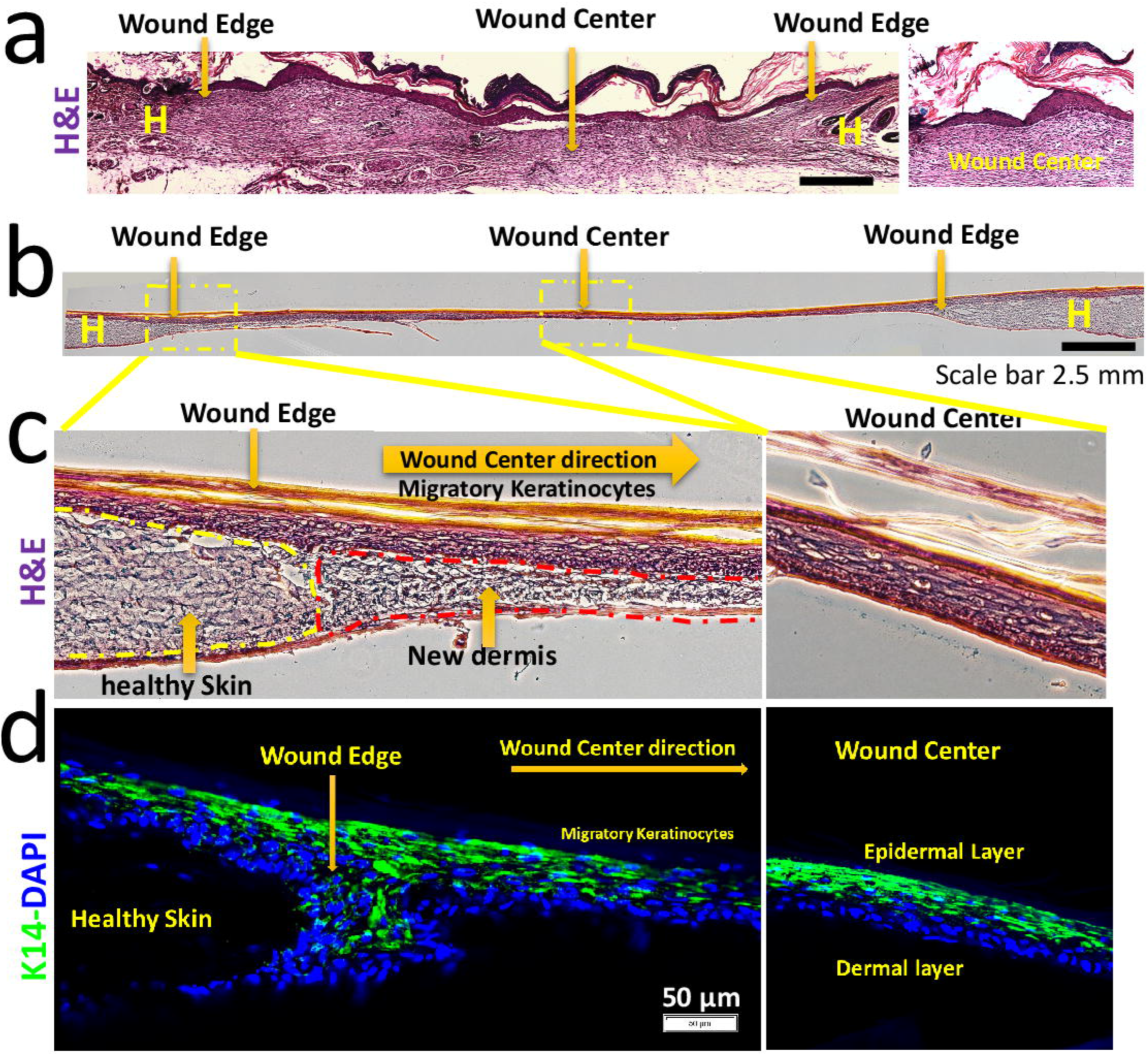
Microscopic evaluation of the wound healing process. Representative images of wound sections on day 10 post-injury with hematoxylin–eosin staining. A full-view reconstruction approach was used for section analysis (a–b). Animal wound sections (a) and skin equivalent wound sections (b). 10 Days after fibrin matrix extrusion, keratinocytes migrated from the healthy skin to the new dermis, reaching the wound center (c–d). Cytokeratin staining for keratinocytes (green) and DAPI-labeling nuclei (blue). Immunofluorescent sections showed keratinocyte migration from the healthy skin (d, left) to the new dermis (d, right). The yellow “H” in the figure indicates healthy skin.

### Bioprinted HSE mimics epidermal differentiation of the animal healing process after injury

Microscopic evaluation of the HSE showed that keratinocytes within the wound center differentiate. After 10 days post wounding, a new stratum corneum was formed (**Figure 4c**, right) in the wound center of the HSE. A similar differentiation process was observed in the wound center of the animal model (**Figure 4a**, right).

## Discussion

Our study demonstrates *in vitro* a 3-D wound healing model incorporating a bioprinted human skin equivalent, a mechanical injury of that skin facsimile, and an FPM–macrophage installed that seals the wound. Importantly, the FPM utilizes the intimate contact of fibrin, pFN, and macrophages to sustain macrophage viability that normally includes the catalysis of M1 to M2 trans-differentiation. This FPM-viable-macrophage mixture was stable for > 7 days and installed as a “bioink” using an extrusion system. Once installed the FPM–macrophage mixture served to guide keratinocytes for emulating the wound closure portion of the mammalian skin repair process (15, 18–22). For com parison, we used a mouse wound model with corticosteroid treatment (**Figure 5a**) that accelerates keratinocyte migration initiation. For example, on day 1 after mouse dermal injury, the fibrin clot structure presented a high infiltration of immune cells (**Figure 5b, c**) as previously shown (5) and a premature initiation of keratinocyte migration after 24 hours. The FPM-macrophage sealed HSE wound seeks to establish a structural and cell co-culture signaling platform for normal keratinocyte migration. This contrasts previous *in vitro* wound healing models that lack a FPM, (19–22) fibroblast incorporation into a dermal layer, (18) and a corresponding epidermal layer (23). Other scaffolds employed in past models were made from non-fibrin nanofibers, (24) collagen– glycosaminoglycan, commercially available collagen matrices, or collagen–elastin materials (25).

**Figure 5.**
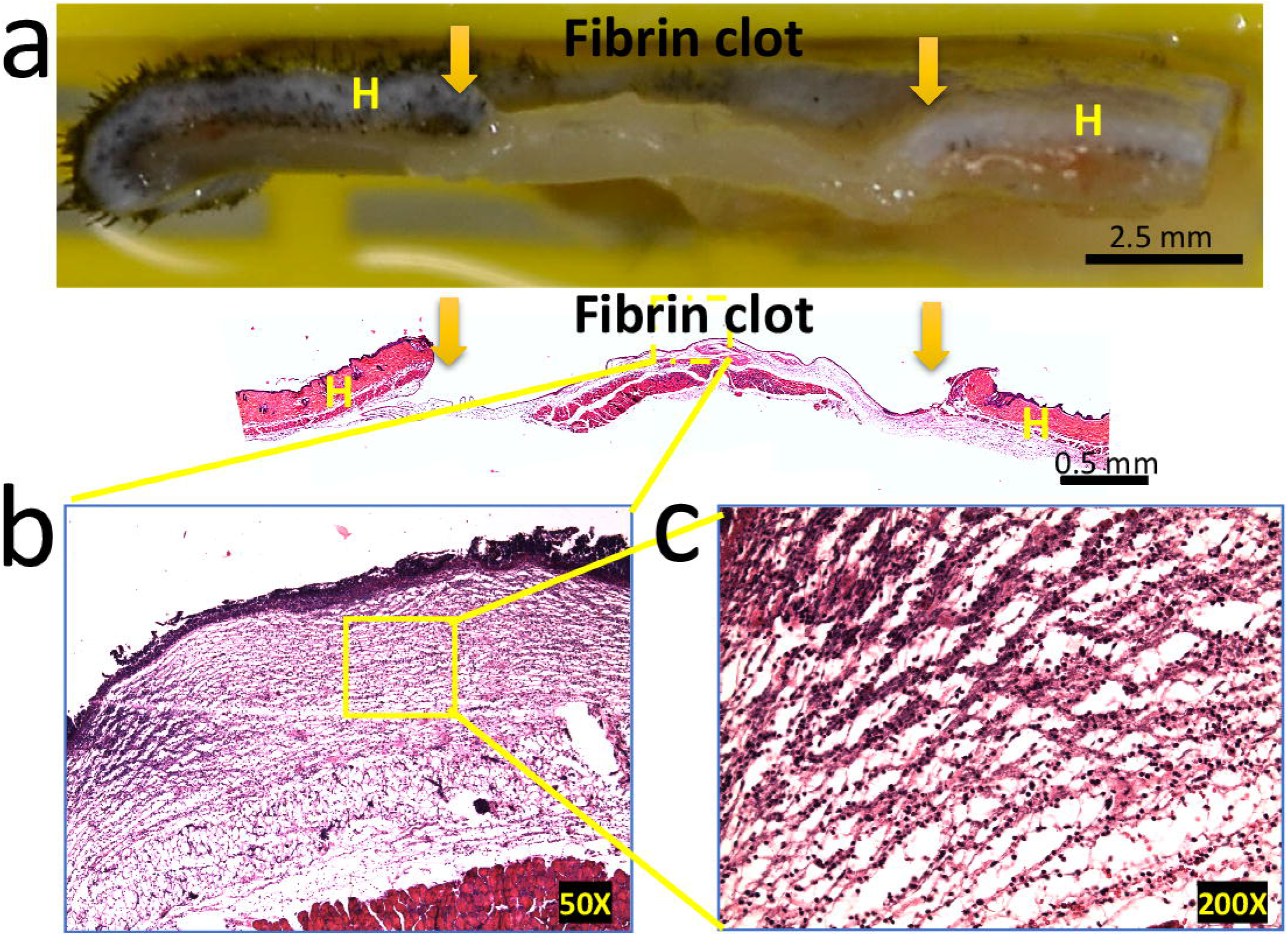
Macroscopic and microscopic evaluation of the fibrin clot structure. Representative macrophotograph of the animal fibrin clot on the top (a) and full view reconstruction on bottom (b). Yellow arrows indicate wound edges, and the yellow “H” indicates healthy skin. Representative fibrin clot sections on day 1 post-injury using hematoxylin–eosin staining, showing immune cell infiltration into the fibrin clot (b–c).

Normal FPM integrin engagement (26, 27) uses Ita5, α5β1, or other fibronectin alpha class receptors belonging to the (RGD)-binding class integrins. These integrins bind to extracellular matrix proteins, such as fibronectin and fibrinogen (28), and are expressed in keratinocytes and fibroblasts. The α5 integrin class is expressed by keratinocytes only after a skin injury and at the wound edge on day 1 post wounding and then steadily progresses to decreased levels during migration (2). Previous *in vitro* cell cultures have used supraphysiological concentrations of F1 without fibrin suspension of macrophages (15). Our FPM formulation used a physiologic 10:1 ratio of purified F1 and pFN with a suspension of macrophages to promote keratinocyte wound surface migration. Future formulations will use pFN nanostructures incorporated into the fibrin network.

As a 3D co-culture model, we purposely altered the culture media environment to emulate the progression from an inflammatory to a proliferative healing phase environment. **Table 3** shows the transition of this culture media achieved by sequential media dilution over day 3 to day 10 of HSE healing. For example, we decreased the insulin and FBS concentrations during the wound inflammatory phase and increased the concentration during the proliferative phase. The gradual dilution of the initial inflammatory media by proliferative media every 2.5 days was designed to taper the cytokines and microRNA levels, as well as temporally changed the growth factor signaling produced by the HSE. The need for an inflammatory phase and a transitional dilution series of media typical of inflammation to one typical of proliferation has been documented. In the normal wound healing process, keratinocytes migrate to the wound center at the end of the inflammatory phase (1, 2). Furthermore, *in vitro* models typically show keratinocyte migration immediately to the mimetic wound center after injury (15). Importantly, the re-epithelialization phenomena over the HSE, in this case, displayed stratification and differentiation where prior *in vitro* models did not show stratum corneum formation (15).

A healthy wound healing pathway requires macrophages engagement with the FPM (3, 4). Initially M1 phenotype macrophages promote an inflammatory response at the wound site. The inflammatory phase that is needed for the process of debridement is normally resolved by M1 apoptosis and with a trans-differentiation component of M1 to M2 macrophages (3, 4). Thereafter fibroblast colonization results in the secretion of collagen based provisional scaffolding to stage angiogenesis by endothelial cells (ECs). In previous reports, we isolated specific fractions of pF1/pFN mixture (29, 30). Also, we demonstrated that specific fractions of pF1/pFN polymeric matrix accelerated wound healing process in healthy mice (31). Here, we analyzed the first level compatibility of FPM-EC bioink formulation where EC viability was observed. Based on the feasibility demonstrated here for FPM-macrophage bioink installation into the HSE wound site, our future work will explore the engineering of a vascularized granulation by EC and fibroblasts co-culture using FPM that are more specialized for EC colonization. Importantly, the current model formulation produced no fibrotic scar tissue structures where future studies will strive to develop capillary structures within a sub-epithelial granulation. In summary, the 3D bioprinted dermal wound healing model incorporating FPM-macrophage bioink provides an *in vitro* co-culture study platform for staging re-epithelialization by keratinocytes. These future studies of an analogous subepithelial healing pathway is envisioned using co-culture of keratinocytes, fibroblasts, endothelial cells, and M2 macrophages.

## Innovation

The FPM here is analogous to a tissue sealant composed of a physiologic mixture of macrophage/fibrinogen/fibronectin that is applied to a mechanically wounded, bioprinted dermal tissue culture. This treatment is shown to temporally synchronize a normal re-epithelization process by keratinocyte migration from the surrounding tissue and over the top of the FPM. Both the keratinocyte migration and wound closure phenomena were demonstrated after the treatment by the FPM-macrophage tissue sealant mixture.

## Key Findings

- Human skin equivalents can emulate animal re-epithelization
- Fibrin clot formation using physiological concentrations of human fibrinogen, human fibronectin and human macrophages can be extruded as a “bioink” into a wound facsimile of a bioprinted tissue.
- A transitional culture medium treatment sequence is used to emulate the environment of the normal wound healing process.

## Acknowledgments and funding sources

The review and wound healing research in the authors’ laboratory were supported by Coordination for the Improvement of Higher Education Personnel (88882.434714/2019-01 and 88881.190035/2018-01). This study was financed in part by the Coordenação de Aperfeiçoamento de Pessoal de Nível Superior, Brasil (CAPES), Finance Code 001 and CNPQ 203415/2014-0.

## Author disclosure and ghostwriting

No competing financial interests exist. No ghostwriters were used to write this article.

## About the Authors

**Carlos P. Jara** is a clinical and research nurse, master’s in health institution management, Master in Health Science, and PhD Student in Health Sciences at Faculty of Nursing in State University of Campinas. His expertise are skin repair and healing process, both in vitro and in animal models.

**Carolina Motter Catarino**. Bioprocess and Biotechnology Engineer. Master’s degree in Pharmacy at the University of São Paulo, Brazil. Carolina research with 3D bioprinting of human skin for cosmetic testing. She completed her doctorate in Chemical & Biological Engineering at the Rensselaer Polytechnic Institute (USA).

**Lício Augusto Velloso**. MD obtained from the University of Campinas, PhD from Uppsala University, Sweden. Pos-doc fellow at Campinas University and Harvard University. Currently Full Professor of Medicine, University of Campinas. Former member of the Editorial Board of Clinical Endocrinology. Current member of the Editorial Board International Journal of Obesity.

**Yuguo Lei** is an Associated professor at Nebraska University – Lincoln. PhD at California Institute for Quantitative Biosciences, UC Berkeley, Chemical and Biomolecular Engineering, UCLA MS, Molecular and Medical Pharmacology, UCLA School of Medicine MPhil, Polymer Science, Hong Kong University of Science and Technology BS, Chemistry, Peking University

**William H. Velander** is a Biochemist with a PhD in Chemical Engineering (The Pennsylvania State University, USA) and is a Full Professor at University of Nebraska–Lincoln, USA. He is a pioneer in novel plasma-derived and genetically engineered formulations of human fibrinogen made from the milk of transgenic animals. Professor Velander is an elected fellow of the American Institute of Medical & Biological Engineering.

**Pankaj Karande** is Chemical Engineer with Ph.D., in Chemical Engineering, University of California, USA. Prof. Karande is an Assistant Professor in the Chemical and Biological Engineering Department at the Rensselaer Polytechnic Institute, USA. Prof. Karande’s research program is focused on engineering peptides as novel drugs, drug carriers, affinity agents and multifunctional biomaterials for medical applications. Peptides play vital roles in various biological functions including membrane assembly, cell regulation and immunity. Inspired by their roles in physiological processes, the Karande Lab is evaluating the potential of short peptide sequences as therapeutics for cancer, neurodegenerative diseases, immune disorders and as sub-unit vaccines against infectious diseases.

**Eliana P. de Araujo, MSC, PhD**, is an Associate Professor at Nursing School, and research of the Obesity and Comorbidities Research Center, University of Campinas, Brazil. Her research focuses on molecular and cellular mechanisms of tissue repair and regeneration and mechanisms involved in the central and peripheral inflammation mediated by fatty acids.

## Abbreviations and Acronyms

FPM: Fibrin Provisional Matrix
HSE: Human Skin Equivalent
pFN: Plasma Fibronectin
HUVEC: Human Endothelial Cells
IMDM: Iscove’s Modified Dulbecco’s Medium
HEPES BSS: HEPES Buffered Saline Solution
PFA: Paraformaldehido
EC: Endothelial Cells

